# The Shiga toxin (Stx)-Phage Encoded Ribosomal RNA Methyltransferase Regulates Stx-producing *Escherichia coli* (STEC) Virulence by Blocking Stx-Mediated Inactivation of Bacterial Ribosomes

**DOI:** 10.1101/2023.09.20.558713

**Authors:** Chen Gong, Gerald B. Koudelka

**Affiliations:** Department of Biological Sciences University at Buffalo, Buffalo, NY 14260

## Abstract

Shiga toxin (Stx) produced and released after induction of Stx-encoding prophage resident within Shiga toxin producing *E. coli* (STEC) causes life-threatening illness. We previously identified that a two-subunit Stx prophage-encoded 16S rRNA methyltransferase, M.ECPA8_3172P-PNB-2, which is both uniquely encoded by and commonly found in Stx2- encoding bacteriophage, regulates both prophage spontaneous induction and STEC virulence. We found here that sequential deletion of these two subunits leads to concomitant, progressive reduction in both prophage spontaneous induction and STEC virulence. This observation indicates that these outcomes are linked. The translation activity of extracts made from a Δ*M.ECPA8_3172P*Δ*PNB*-2 Stx prophage-containing strain was lower that of extracts made from either the methyltransferase replete STEC strain or from a strain that did not contain a Stx-encoding prophage. We found that the Δ*M.ECPA8_3172P*Δ*PNB*-2 STEC strain contained significantly fewer ribosomes that did the methyltransferase replete STEC strain. These observations suggested that the M.ECPA8_3172P-PNB-2 methyltransferase may block Stx-mediated ribosome inactivation. Consistent with this idea, we found that translation extracts made from STEC expressing M.ECPA8_3172P-PNB-2 are more resistant to Stx-mediated inactivation than are those made from Δ*M.ECPA8_3172P*Δ*PNB*-2 STEC. These findings indicate the M.ECPA8_3172P-PNB-2 methylation of 16S rRNA protects the ribosome from Stx-mediated inactivation, thereby allowing more phage and more Stx to be spontaneously produced. Direct 16S rRNA sequencing identified 4 putative M.ECPA8_3172P-PNB-2 methylation sites, all of which map onto the RNA polymerase contacting surface of the 30S ribosome subunit in the expressome, suggesting the M.ECPA8_3172P-PNB-2 may protect the ribosome from inactivation by stabilizing this complex.

## Introduction

Shiga toxin-producing *E. coli* (STEC) cause life-threatening diseases by synthesizing and releasing Shiga toxin (Stx). All STEC strains contain one or more prophage which can encode either of the two different Stx types: Stx1 or Stx2 (*1*). Human intoxication with STEC can induce the symptoms of life-threatening hemolytic uremic syndrome (HUS) (*2, 3*). Stx is a ribosome inactivating toxin. It removes an adenine from the sarcin-ricin loop of the large ribosomal subunit rRNA (*4*), thereby inhibiting protein synthesis in target cells (*5–7*). This damage triggers cellular apoptosis, even a single Stx molecule may be enough to kill a cell (*8, 9*). The sarcin-ricin loop is conserved between bacteria and eukaryotes (*10, 11*), thus ribosomes of both bacterial and eukaryotic cells are vulnerable to the Stx attack (*12, 13*).

All Stx-encoding phages are temperate phages. Upon infection, these phages can either replicate and destroy the host bacteria (lytic cycle) or enter into a dormant state called lysogeny, in which the viral genome carrying the Stx encoding genes is integrated into the genome of host cell. STEC infection of humans occurs through the ingestion contaminated food or water. The prophage in these strains are in the lysogenic state. The phage genome can remain dormant for an extended period (*14–16*). The initiation of Stx-mediated disease requires that the prophage in STEC strains switch from lysogenic to lytic growth (*14–16*).

During lytic growth, the phage genome excises from the host chromosome, is packed into viral particles and these released from the cell upon lysis. Lytic growth of these phage also results in the activation phage late genes, including *stx1* and/or *stx2*. The stx2 genes are under the control of promoters that are active exclusively during the later stages of lytic growth. Thus Stx2 is only produced during phage lytic growth. The *stx1* genes have an additional promoter that is activated under iron-limiting conditions (*17, 18*), meaning Stx1 can be produced also in the absence of prophage induction. Regardless, the Stx protein cannot be actively transported out the bacterial cell. Moreover high level production of both Stx 1 and Stx only occurs during lytic growth. Therefore, Stx production and its subsequent release requires lytic growth of the prophage (*19–22*). Thus lytic growth of the phage and consequent lysis of the bacterial host are essential to STEC pathogenesis, irrespective of Stx type.

The switch from lysogenic to lytic growth is called induction. The amount of Stx produced is proportional to the number of bacteria that undergo induction (*23, 24*). Strains that produce higher levels of Stx are thought to be more virulent to humans. Thus, the mechanisms that control induction are crucial to the initiation and progression of STEC- mediated disease.

Induction of Stx prophage requires the removal of the phage-encoded *c*I repressor protein from its DNA binding sites. Prophage induction can occur spontaneously, without any external stimuli (*25*), or it can also be induced by various environmental factors such as UV radiation, DNA-damaging chemicals, host cell stressors and other factors (*23, 24*). Several investigations show that prophages resident in highly virulent STEC spontaneously induce at high frequency (*17, 26–28*). The high frequency of spontaneous induction promotes the development of HUS during STEC infection (*25, 29–31*).

Earlier studies proposed that prophage spontaneous induction occurred resulted from stochastic variations in gene expression (*32–34*). However, recent work shows that high spontaneous induction frequencies of temperate prophages increases virulence, promotes biofilm formation and improves the spreading of genes through horizontal gene transfer (*25, 29, 35*). These findings suggest selection for traits that allow higher spontaneous induction frequencies could be part of strategy that increases phage fitness (*25*).

Our earlier work indicated that the increased spontaneous induction frequencies of Stx-encoding prophage resulted in part from the lower intracellular levels of their *c*I repressor found in lysogens containing these phages than in lysogens that harbor non-Stx encoding lambdoid phage (*36, 37*). Consequently, less repressor is available to occupy its binding sites in Stx prophage lysogens, increasing the chances for the prophage to switch to lytic growth as compared to other lambdoid prophages (*27, 28*). However, it is clear that other prophage functions also play a role in promoting spontaneous induction of Stx-encoding prophage.

We recently identified a methyltransferase gene that is found uniquely in Stx-producing phage and is especially common in phages that harbor genes that produce Stx2 isoform. Disruption of this gene decreases prophage spontaneous induction frequency and reduces the ability of STEC to kill eukaryotic cells (Chapter 2). Initial annotation identified the function of this gene as an adenine DNA methyltransferase. Yet, *in vitro* study showed that the purified methyltransferase (M.ECPA8_3172P), displays only non-specific methyltransferase activity with DNA or RNA substrates. However, the methyltransferase enzyme is co-expressed and co-purifies with another subunit, called PNB-2 and the purified holoenzyme preferentially methylates 16 rRNA.

To our knowledge the M.ECPA8_3172P-PNB-2 holoenzyme comprises the first known phage-encoded RNA methyltransferase. However, it is unclear how methylating 16s rRNA could affect prophage spontaneous induction and eukaryotic cell killing. To better understand how the M.ECPA8_3172P-PNB-2 holoenzyme affects this process, we created a series of STEC strains, each of which bears a deletion of the genes encoding either M.ECPA8_3172P, PNB-2 or both and examined the impact of these changes on eukaryotic cell killing, prophage spontaneous induction, translation and ribosome stability. Here, we show that the methylation performed by M.ECPA8_3172P-PNB-2 apparently affects cell killing and prophage spontaneous induction by enhancing translation by an increasing ribosome viability in Stx- encoding strains. Our data indicate these effects are mediated by methylation-dependent inhibition of Stx-dependent bacterial ribosome inactivation. These findings suggest that the phage actively protects the host cell translation machinery in order to increase both amount of Stx released and the number of phages produced.

## Results

### Both the M.ECPA8_3172P and PNB-2 contribute to induction regulation and eukaryotic cell killing

We showed previously that deletion of a gene encoding a subunit of a 16s rRNA methyltransferase from Stx-encoding prophage decreases both the prophage spontaneous induction frequency of the prophage and the virulence of bacteria bearing the prophage. High *in vitro* activity and specificity of the methyltransferase requires a holoenzyme that consists of two subunits, the methyltransferase subunit, M.ECPA8_3172P, and an RNA recognizing subunit, PNB-2. BLAST searches indicate that the methyltransferase gene is only present in bacteria bearing Stx-encoding prophage. However not all these phages/strains contain the gene encoding PNB-2. Given that most of the variation in STEC virulence is attributed to the specific prophage(s) resident in the strain, we to explore how the presence of the M.ECPA8_3172P subunit alone or the absence of both M.ECPA8_3172P and PNB-2 affects prophage growth and bacterial virulence.

For this investigation, we first deleted *gp6* gene encoding PNB-2 from ϕPA8, creating the ϕPA8Δ*PNB-*2, which thereby expresses only M.ECPA8_3172P. Subsequently we also deleted *gp5* gene which encodes M.ECPA8_3172P, creating the double knock-out phage, ϕPA8Δ*M.ECPA8_3172P*Δ*PNB-2* (Figure 1). We formed lysogens with these phages in *E. coli* strain MG1655. Since these genes are part of a larger operon (P. Berger, personal communication), prior to further analysis, we first confirmed that these deletions did not affect transcription of downstream genes in the operon (Figure S1).

**Figure 1.**
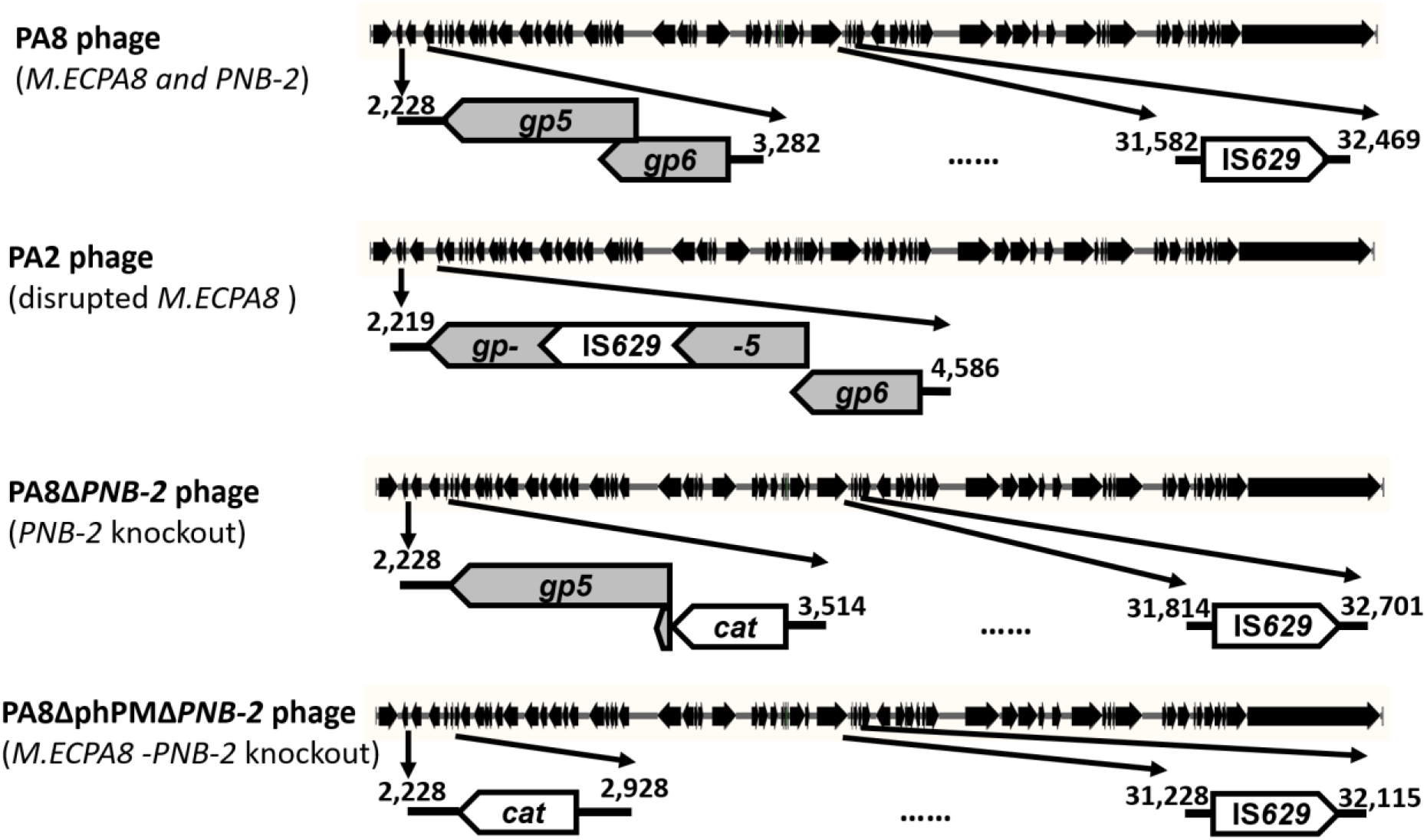
The genomic structures of Stx-encoding phages used in this study. Shown is a schematic representation of the genomes of ϕPA8 highlighting the positions of *gp5* and *gp6* gene that encodes the M.ECPA8_3172P and PNB-2 and the IS*629* mobile element inserted into a non-coding region; ϕPA2 highlighting the disruption of *gp5* by the IS*629* insertion element; ϕPA8Δ*PNB-*2 in which we replaced the *gp6* gene with that for chloramphenicol acetyl transferase (*cat*); and ϕPA8Δ*M.ECPA8_3172P*Δ*PNB-2* (bottom) in which we replaced both the genes of M.ECPA8_3172P and PNB-2 with the *cat* gene.

We compared spontaneous induction frequency of the prophage ϕPA8Δ*PNB-2*, ϕPA8Δ*M.ECPA8_3172P*Δ*PNB-2* with that of ϕPA8, which expresses both M.ECPA8_3172P and PNB-2 and ϕPA2, in which the gene of M.ECPA8_3172P is disrupted by an IS*629* element and thus expresses only PNB-2. We found that prophages expressing only one of the two genes that comprise the methyltransferase holoenzyme have a lower spontaneous frequency than does ϕPA8, which expresses both subunits (Figure 2A, black vs grey circle and triangle). This trend becomes particularly obvious after 90 min of growth (P < 0.01) at which time spontaneous phage production by MG1655::ϕPA2 and MG1655::ϕPA8Δ*PNB-2* strains has essentially stopped, while MG1655::ϕPA8 continues to release phage at an accelerating frequency. Deleting both of the genes that encode the methyltransferase holoenzyme essentially eliminates spontaneous induction (Figure 2A, white diamonds); at virtually all time points (P < 0.01) the MG1655::ϕPA8Δ*M.ECPA8_3172P*Δ*PNB-2* produced significantly less phages than any of the other strains. These findings confirm that both M.ECPA8_3172P and PNB-2 subunits have a role in regulating prophage spontaneous induction.

**Figure 2.**
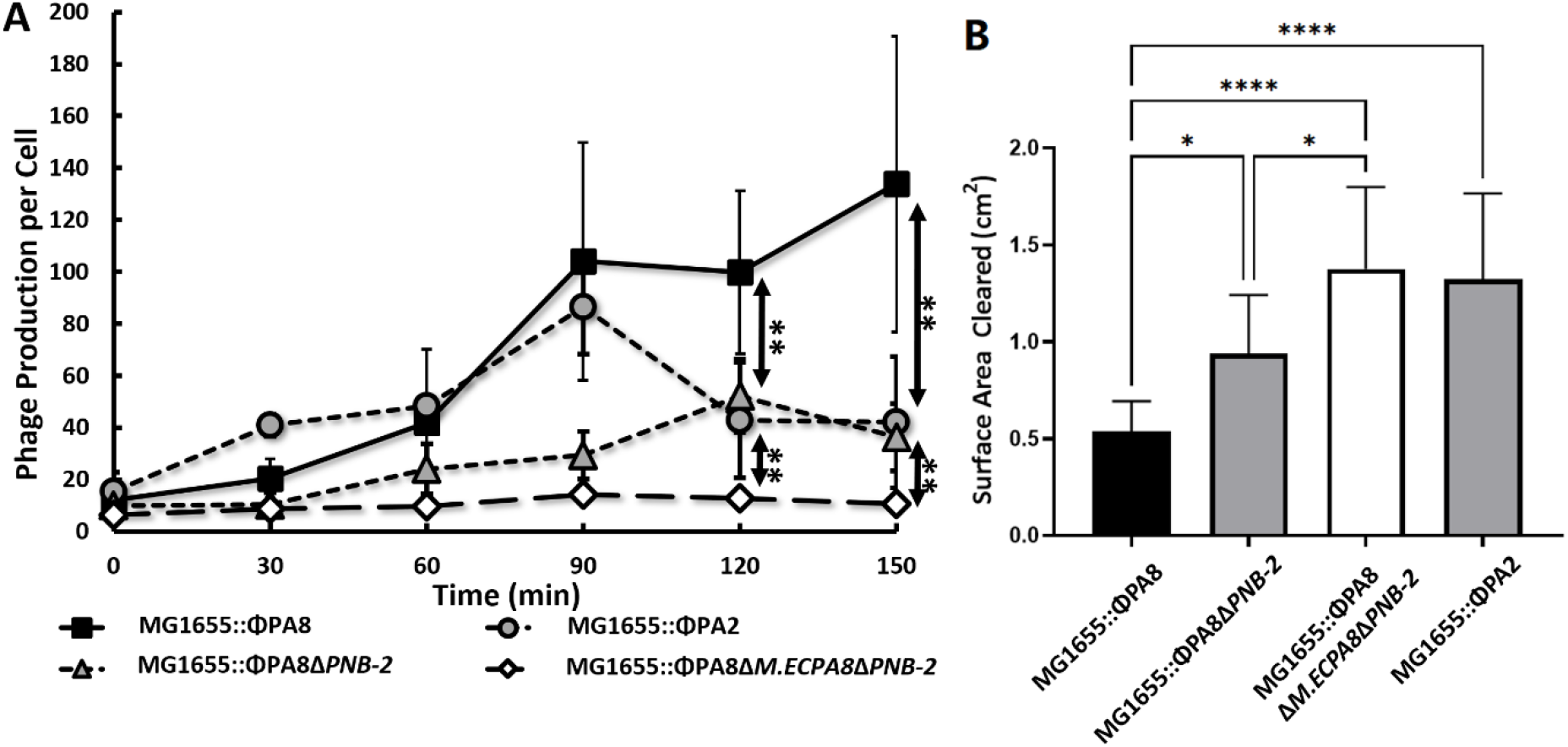
The spontaneous induction frequency and amoeba killing ability of *E. coli* MG1655 strain bearing different Stx-encoding prophages. A. Overnight cultures of different lysogen strains were diluted 40-fold and grown for 2.5h. Subsequently the number of *E coli.* cells and free phages were measured by qPCR every 30 min as described in Methods and Materials. Shown is the number of phages produced per *E. coli*. Error bars represent standard deviations and each data point was determined from at least 3 biological replicates and each biological replicate was measured from at least 3 technical replicates. ** *P* < 0.01; * *P* < 0.05. **B.** The effect of expressing *M.ECPA8_3172P, and/or PNB-2* on *A. castellanii* predation of *E. coli. A. castellanii* (10^6^ cells in a volume of 10μl) were separately spotted on PPG plates seeded with 10^8^ cells of *E. coli* strains MG1655::ϕPA8, MG1655::ϕPA8Δ*PNB-2*, MG1655::*ϕPA8ΔM.ECPA8_3172PΔPNB-2* or MG1655::ϕPA2. *A. castellanii* were allowed to grow on these plates for 5 days at 28.5℃. If *A. castellanii* graze the bacteria on plate, it will form a plaque on the bacterial lawn. The size of the plaques was determined using ImageJ. Error bars represent standard deviations from at least 3 independent quadruplicate measurements. The one-way analysis of variance (ANOVA) followed by a post-hoc implementation of Tukey’s multiple comparisons correction. Comparisons were made using the adjusted P value to determine the statistical significance of pairwise comparisons. ****: P < 0.0001; * *P* < 0.05.

We wished to examine whether the deletions of the individual subunits of the phage-encoded methyltransferase also affected the ability of *E. coli* bearing Stx-encoding prophage to kill eukaryotic cells. For this experiment, we used a solid phase predation assay (Methods and Materials, see also (*38*)) to examine the ability of the eukaryotic predator amoeba, *Acanthamoeba castellanii* (*A. castellanii*) to grow on the strains bearing deletions of the gene encoding individual subunit or both subunits. For these experiments, the four different toxin-encoding *E. coli* strains: MG1655::ϕPA8, MG1655::ϕPA2 and MG1655::ϕPA8Δ*PNB-2* and MG1655::ϕPA8Δ*M.ECPA8_3172P*Δ*PNB-2* were separately seeded onto PPG agar plates (see Methods and Materials) and incubated at 37°C for 1 h. Subsequently, amoebae were spotted on the center of the plate, sealed with parafilm and incubated at 28.5°C for 5 days. After incubation, the surface area cleared by the amoebae on each plate was measured and recorded. We showed previously that the size of amoebae plaque formed on toxin-encoding *E. coli* is inversely proportional to the amount of Stx-mediated killing (*39*).

Amoebae consumed all four bacterial strains, forming plaques in the bacterial lawns (Figure 3B). However, the plaques formed on MG1655::ϕPA8 were much smaller than those formed on the other strains (Figure 3B, black bar). Consistent with our previous findings, this result indicates that Stx produced by this strain efficiently kills *A. castellanii*. Amoebae formed significantly larger plaques on the strain bearing deletion of the RNA binding subunit, MG1655::ϕPA8Δ*PNB-2*, indicating this strain does not kill amoeba as effectively as the strain encoding both methyltransferase subunits (Figure 3B, grey bar). Amoebae formed even larger plaques on the strain bearing deletions of both the RNA binding and methyltransferase subunit MG1655::ϕPA8Δ*M.ECPA8_3172P*Δ*PNB-2* (Figure 3B, white bar). The plaques formed on this strain were similar in size to those formed on MG1655::ϕPA2, the strain that bears a disruption of gene encoding the methyltransferase subunit M.ECPA8_3172P (Figure 3B, grey bar). We note that the effect of the deletions on amoeba toxicity substantially mirrors that of their effect on spontaneous induction frequency (compare Figure 3A & B). This observation suggests that M.ECPA8_3172P and/or PNB-2 affects the ability of the STEC strain to killing eukaryotic cells by regulating spontaneous induction frequency.

**Figure 3.**
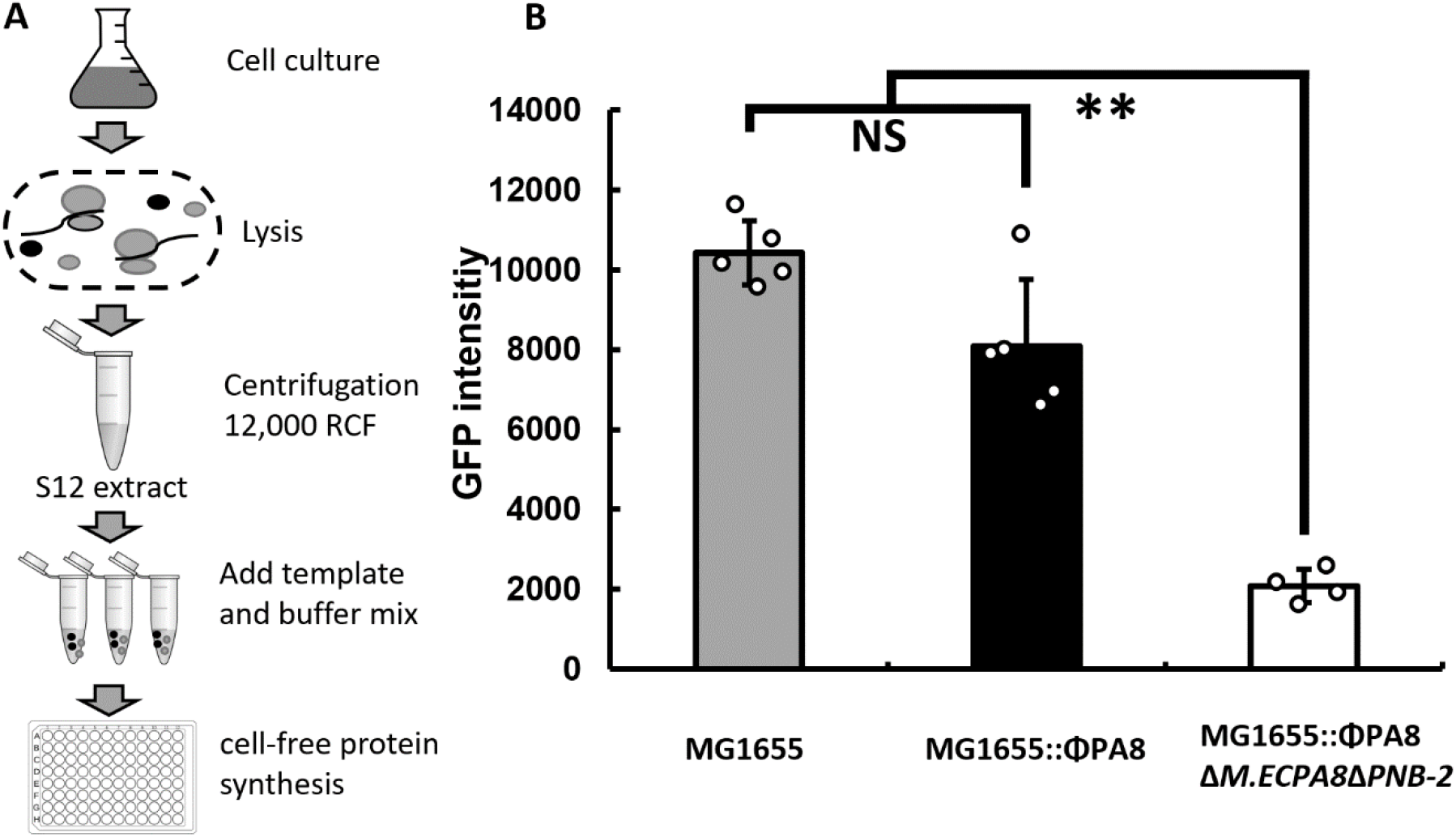
M.ECPA8_3172P-PNB-2 increases overall translation efficiency of the *E coli.* strain bearing Stx-encoding prophage. A. Schematic representation of cell extract preparation by S12 procedure; **B.** Cell-free protein synthesis using the plasmid bearing *sfGFP* gene (p-*GFP*) as template with S12 extracts made from *E. coli* strains bearing ϕPA8, ϕPA8Δ*M.ECPA8_3172P*Δ*PNB-2* or no prophage. The GFP intensity was measured as described in Methods and Materials. Error bars represent standard deviations and each data point was determined from at least 3 biological replicates and each biological replicate was measured from at least 3 technical replicates. ** *P* < 0.01; **NS** not significant *P* > 0.05.

### M.ECPA8_3172P-PNB-2 regulates overall translation efficiency in present of prophage

Our previous findings showed that in *E. coli* the M.ECPA8_3172P-PNB-2 holoenzyme methylates 16s ribosomal RNA. Most of the known modifications introduced into 16s rRNA by host-encoded methyltransferases play a role in regulating translation initiation and/or fidelity (*40, 41*). Since the regulation of translation may affect prophage and subsequent Stx production during spontaneous induction, we determined whether the M.ECPA8_3172P- PNB-2 holoenzyme regulates translation.

Bacterial cells lyse as a consequence of prophage induction. This condition limits our ability to straightforwardly examine the effect of M.ECPA8_3172P-PNB-2 on translation *in vivo*. Consequently, we developed a cell-free protein synthesis system (CFPS) to probe the effect of M.ECPA8_3172P-PNB-2 methylation on translation (Figure 3A, see also Methods and Materials). Briefly we monitored the production of green fluorescence protein (GFP) in S12 extracts (*42*) prepared from *E. coli* strains bearing Stx-encoding prophages that express neither or both the M.ECPA8_3172P-PNB-2 subunits.

We found that the overall translation efficiency of GFP in extracts made from lysogens bearing a Stx-encoding prophage containing the methyltransferase genes is similar to that of the prophage-free host strain (Figure 3B, black bar vs grey bar). In contrast, the amount of GFP produced in extracts prepared from the MG1655::ϕPA8Δ*M.ECPA8_3172P-*Δ*PNB-2* strain is significantly lower than that produced in extracts prepared from either MG1655 or MG1655::ϕPA8 (Figure 3B, white bar). These findings suggest that the presence of Stx-encoding prophage decrease translation in *E. coli* strain during spontaneous induction, while the 16s ribosome methylation performed by M.ECPA8_3172P-PNB-2 holoenzyme rescues this decrease.

### M.ECPA8_3172P-PNB-2 increases ribosome abundance in presence of Stx-encoding prophage

The results in Figure 3 clearly show that the M.ECPA8_3172P-PNB-2 holoenzyme blocks the negative impact of Stx-encoding prophage carriage on translation. This observation suggests that methylation overcomes the impact of a Stx prophage-encoded translation “inhibitor”. We hypothesize this inhibitor may affect ribosome number (*43–45*), ribosome efficiency (*46*), or both. To begin to distinguish between these possibilities, we measured the amount of 16s rRNA by RT-qPCR as a proxy to monitor the number of ribosomes that accumulate over time in *E. coli* strains bearing Stx-encoding prophage that did or did not express the M.ECPA8_3172P-PNB-2 holoenzyme. The measured amount of 16s rRNA in these strains was normalized to the amount of the mRNA transcribed from the housekeeping *uidA* gene.

We found a significant difference in ribosome abundance among the strains carrying different prophages (Figure 4). At virtually all times after initiating growth, the MG1655 strain carrying wild-type ϕPA8 contained a higher level 16s rRNA (Figure 4, black square) than did the strain bearing deletion of both *M.ECPA8_3172P* and *PNB-2* (Figure 4, white diamonds) This trend became significant after 30 min of growth (P > 0.05). Interestingly, the time course of expression of 16s rRNA mirrored that of the overall translation efficiency (Figure 3) and prophage spontaneous induction (Figure 2). These findings suggest that the M.ECPA8_3172P-PNB-2 holoenzyme regulates ribosome abundance, consequently affecting translation activity and thereby prophage spontaneous induction.

**Figure 4.**
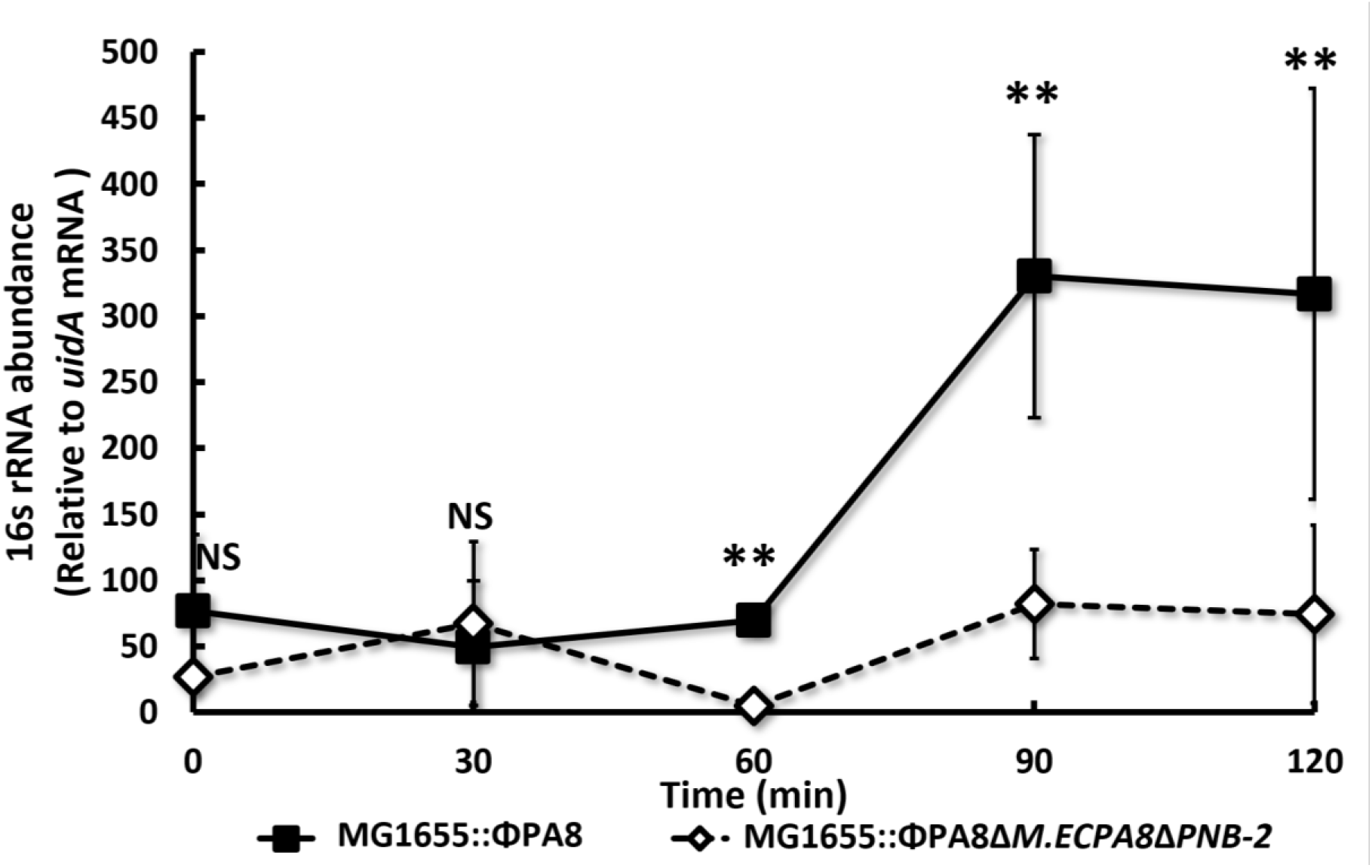
M.ECPA8_3172P-PNB-2 increases ribosome abundance in the *E coli.* strain bearing Stx-encoding prophage. Overnight cultures of MG1655::ϕPA8 and MG1655::ϕPA8Δ*M.ECPA8_3172P*Δ*PNB-2* were diluted 40-fold and grown for 2h. Subsequently the amount of 16s rRNA and *uidA* mRNA were measured by RT-qPCR every 30min as described in Methods and Materials. Shown is the 16s rRNA abundance relative to *uidA* mRNA abundance. Error bars represent standard deviations and each data point was determined from at least 3 biological replicates and each biological replicate was measured from at least 3 technical replicates. **\*\*** *P* < 0.01; **NS** not significant *P* > 0.05.

### M.ECPA8_3172P-PNB-2 protect ribosomes from attack by Stx

Stx is a ribosome inactivating protein that removes a specific adenine in the universally conserved α-sarcin/ricin loop (SRL) of the large rRNA, inhibiting protein synthesis in target cells (*4*). The *M.ECPA8_3172P* gene is commonly found in organisms that also encode Stx2. Analysis of *E. coli* or phages that express the Stx2 isoform indicate the methyltransferase gene is found in 45% of these organisms. Since bacterial 23s rRNA also contains the α- sarcin/ricin loop, *E. coli* ribosomes can be inactivated by Stx (*12, 47*). Knowing this and given the frequency with which Stx2 and the methyltransferase co-occur, the finding that expression of the methyltransferase is “protective” of translation (Figure 3 & 4), suggests that M.ECPA8_3172P-PNB-2 may protect the bacterial ribosomes from Stx inactivation, thereby maintaining their activity and abundance.

To test this idea, we examined the effect of expressing Stx2 in transcription/translation extracts made from strains that do or do not express the M.ECPA8_3172P-PNB-2 methyltransferase on translation of GFP. Briefly in this experiment, we first added the plasmid carrying *stx_2_* gene (p-*stx_2_*) to the S12 extracts prior adding the p-*GFP* and then compared the change of overall translation efficiency between different S12 extracts (see Methods and Materials). The relative change of translation efficiency is calculated as follow:

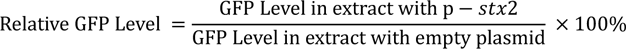

We found adding increasing the amounts of p-*stx_2_* reduced GFP translation in all S12 extracts (Figure 5). The observation is consistent with prior findings and confirms that bacterial ribosomes are inactivated by Stx2. However at all concentrations of p-*stx_2_*, the relative amount of GFP translated in extracts made from MG1655::ϕPA8 was significantly higher than in extracts made from MG1655::ΦPA8Δ*M.ECPA8_3172P*Δ*PNB-2* (Figure5, black bars vs white bars). This finding indicates that M.ECPA8_3172P-PNB-2 methylation provides protection to ribosomes against the Stx2 attack thereby maintaining translation when Stx2 is expressed.

**Figure 5.**
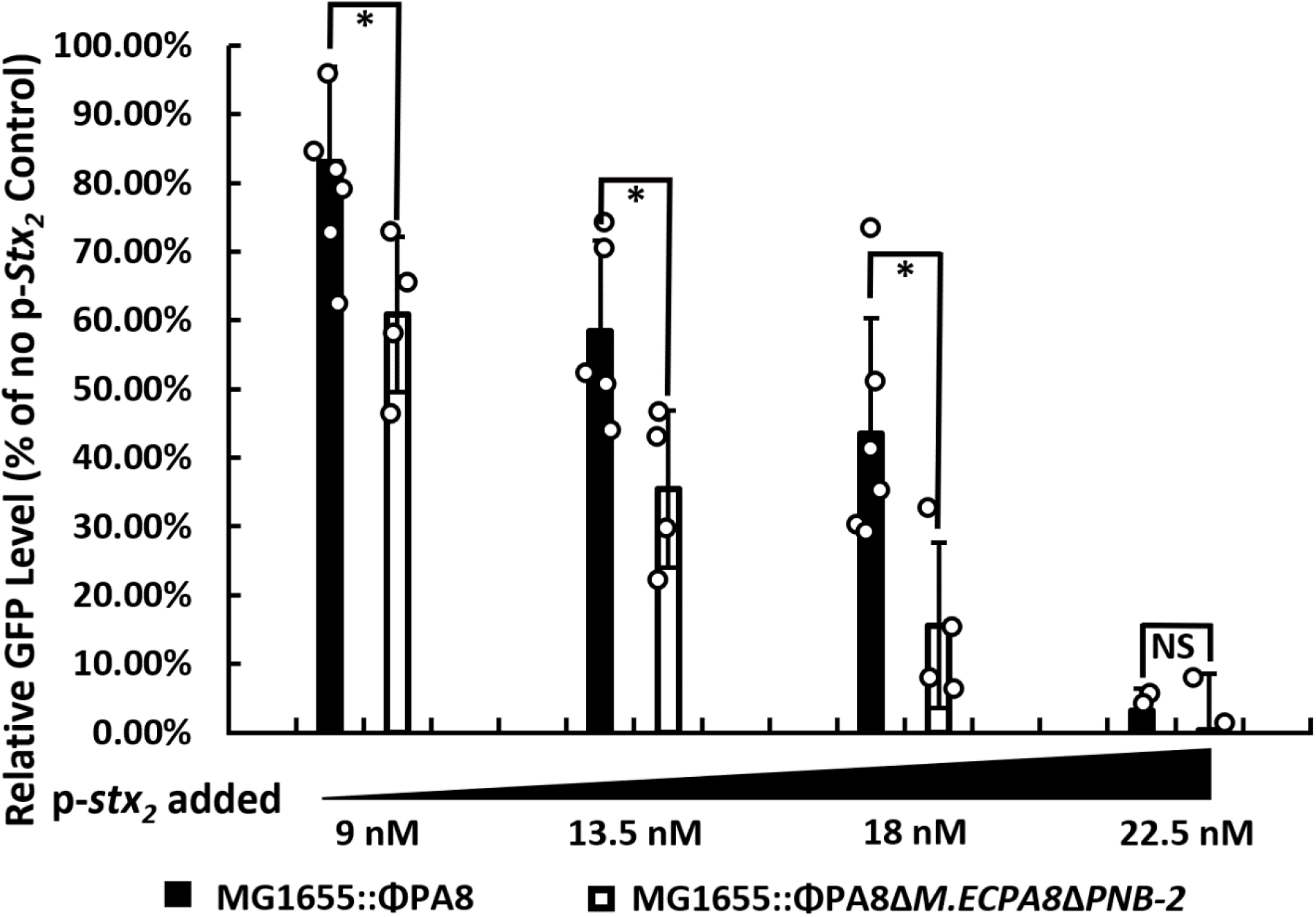
M.ECPA8_3172P-PNB-2 maintains translation activity in presence of Stx2. Show are the results of cell-free protein synthesis using S12 extract made from MG1655::ϕPA8 and MG1655::ϕPA8Δ*M.ECPA8_3172P*Δ*PNB-2* strain in the presence of different amount of the plasmid carrying *stx_2_* gene. The GFP intensity was measured as described in Methods and Materials. Error bars represent standard deviations and each data point was determined from at least 3 biological replicates and each biological replicate was measured from at least 3 technical replicates. **\*** *P* < 0.05; **NS** not significant *P* > 0.05.

### The potential methylation target of M.ECPA8_3172P-PNB-2

To gain insight into the mechanism by which M.ECPA8_3172P-PNB-2 methylation may help maintain ribosome function, we used direct RNA sequencing to identify the base(s) on 16S rRNA that are modified by this enzyme. For this experiment, we prepared and sequenced three samples, 1) an unmodified *in vitro* transcribed control 16S rRNA; 2) 16S rRNA purified from 30S ribosomal subunits isolated from MG1655 (no phage control) and; 3) 16S rRNA purified from 30S ribosomal subunits isolated from MG1655::ϕPA8 (Methods and Materials). Positions of M.ECPA8_3172P-PNB-2 methylation were identified by using xpore (*48*) to perform pairwise comparison signal level data ionic flow features (signal intensity, dwell time, etc.) obtained from the three samples. Our analysis revealed that k-mers containing bases 832, 1098, 1100, 1132 and 1190 in 16S rRNA isolated from MG1655::ϕPA8 showed significantly disturbed electrical signals (log_10_ (p-value) <-100) as compared to those obtained with 16S rRNA isolated from MG1655 (Fig 3-6A). Mapping these positions onto the structure of the 70s ribosome shows that the k-mers encompassing bases at these positions are either mostly or completely solvent exposed on top surface of the “head” of the 30S ribosomal subunit (Figure 3-6B and C). We note that this surface comprises the site that contacts the transcribing RNA polymerase and is thought to play a role in transcription-translation coupling in bacteria (*49–52*) (Figure 3-6D).

**Figure 6.**
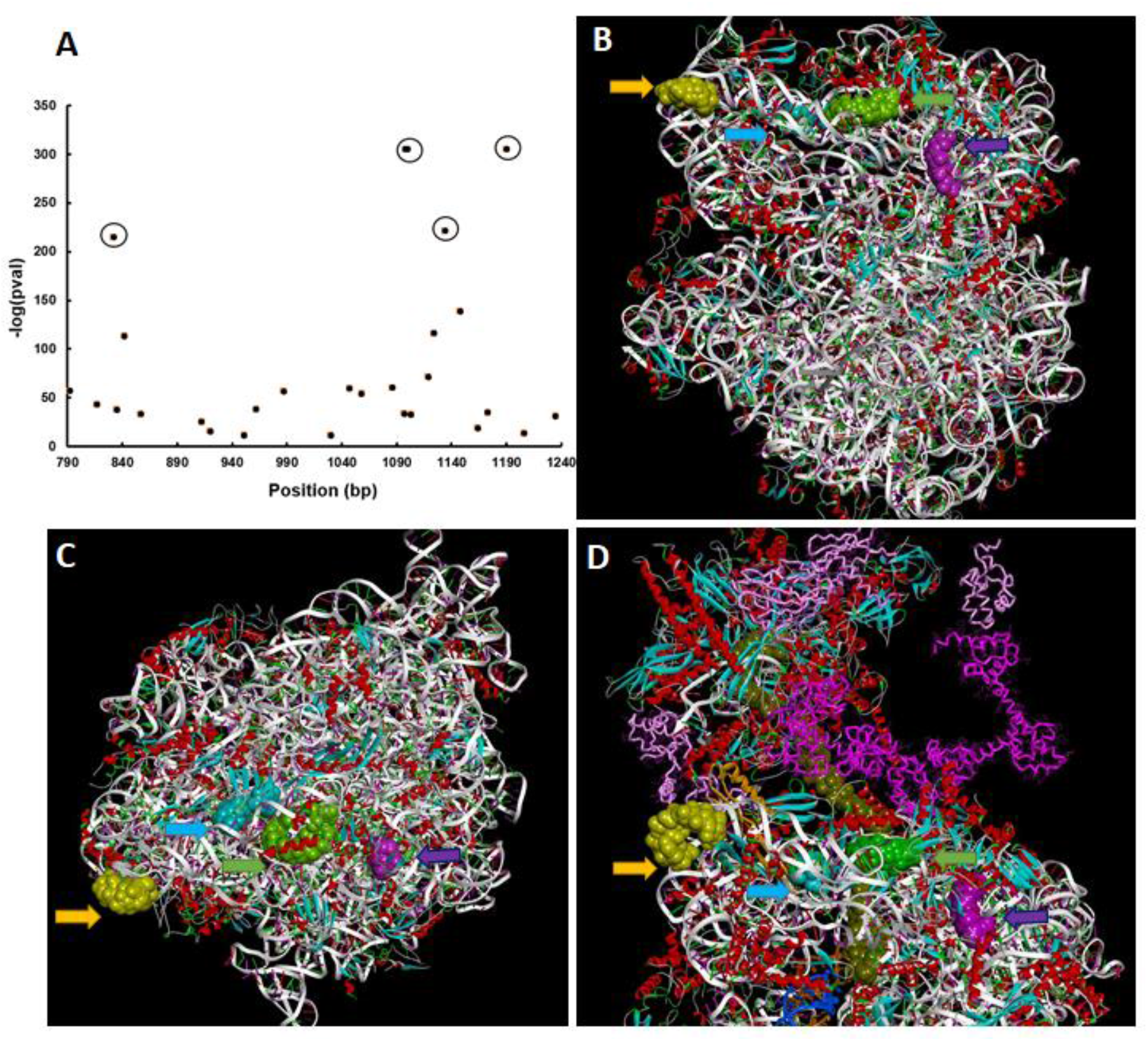
Identity and locations of 16S rRNA bases modified by M.ECPA8_3172P-PNB2. **(A)** Plot of the −log_10_ (pvalue) for the comparison of modification rates between 16S rRNA prepared from MG1655 and MG1655::ϕPA8. Shown is the region of the plot containing *k*-mers who log_10_ (pvalue) is ≤ −150. The circles identify *k*-mers centered at positions 823, 1098 & 1100, 1132 and 1190. **(B)** Location of the bases in the *k*-mers identified in identified in **(A)** mapped onto the structure of the 70S *E. coli* ribosome (*53*). The bases in the *k*-mers centered at the positions in **(A)** are rendered in CPK and differentially colored 832, dark pink; 1098/1100, green; 1132, yellow and; 1190, light blue. **(C)** Representation as in **(B)** rotated 90^◦^ about an axis that bisects the small and large subunits. **(D)** Location of the bases in the k-mers identified in identified in **(A)** mapped onto the structure of the structure of the *E.coli* coupled transcription-translation complex B1 (TTC-B1) that contains an mRNA with a 24 nt long spacer (olive green CPK),and transcription factors NusA (dark magenta tube) and NusG (light magenta tube) The orientation of the 70S ribosome is similar to that shown in **(B)**. Figures B-D were created with BIOVIA, Dassault Systèmes, Discover Studio Visualizer, version 21.1, San Diego: Dassault Systèmes, 2020.

## Discussion

We previously identified a two-subunit Stx prophage-encoded 16S rRNA methyltransferase, M.ECPA8_3172P-PNB-2 that is both uniquely encoded by and commonly found in Stx2-encoding bacteriophage (Chapter 2). This enzyme regulates both the amount of phage spontaneously released by STEC as well as the ability of STEC to kill eukaryotic cells. We show here that that cells that encode both Stx and M.ECPA8_3172P-PNB-2 suggest have increased 16S rRNA levels as compared to Stx-encoding cells lacking the methyltransferase. Similarly, Stx-encoding cells containing the M.ECPA8_3172P-PNB-2 methyltransferase have higher translational activity than do Stx-encoding cells lacking this enzyme. Moreover, the abundance of 16S rRNA mirrors that of both the overall translation efficiency (Figure 3) and amount of phage spontaneously produced by Stx-encoding cells (Figure 2). These finding indicate that the Stx-encoding cells containing the M.ECPA8_3172P-PNB-2 methyltransferase contain more ribosomes than do Stx-encoding cells lacking this enzyme.

We also found that the amount of spontaneously produced phage mirrors the overall virulence of Stx-encoding strain. That is, the increase in translation in STEC that encode the M.ECPA8_3172P-PNB-2 methyltransferase results in an increase in number of phages spontaneously released by Stx-encoding cells as well as the eukaryotic cell killing efficiency of these cells (Figure 2A & B). These findings confirm the long-held hypothesis that the observed the level of prophage spontaneous induction plays a role in regulating bacterial virulence during human infection (*25, 28, 54, 55*). Our results also show that spontaneous induction and consequent Stx production are regulated by the M.ECPA8_3172P-PNB-2 methyltransferase.

Our results indicate that expression of M.ECPA8_3172P-PNB-2 increases translation activity in STEC bearing Stx-encoding prophage by blocking Stx-mediated ribosome inactivation. This observation suggests a model to explain how 16S rRNA methylation regulates the amount of phage that STEC spontaneously produces and ultimately the virulence of these strains (Figure 7). According to our model, the seemingly stochastic increase in transcription of prophage late genes in a given cell results in the production of Stx along with those responsible for phage replication, assembly and release. Since Stx can inactivate bacterial ribosomes, the amount of phage and Stx released by the cell is limited by Stx effects on translation activity. We showed previously that M.ECPA8_3172P-PNB-2 is produced constitutively and our measurements indicate that the entire pool of 16S rRNA has been modified by this enzyme (Chapter 2). Therefore, during the spontaneous initiation of the lytic program, all ribosomes are resistant to Stx inactivation. Consequently, our model suggests that cells containing the M.ECPA8_3172P-PNB-2-modified ribosomes produce more phage and more Stx per cell than do cells that express Stx but not M.ECPA8_3172P- PNB-2 (Figure 7). It is important to note that according to this idea, M.ECPA8_3172P-PNB-2 does not affect the fraction of cells in the population that undergo induction. Instead the increased number of phages released and the increased virulence of M.ECPA8_3172P-PNB- 2-encoding STEC under non-inducing conditions results from each cell undergoing spontaneous induction produce more phage and Stx as a result of methylation dependent resistance of the ribosomes to Stx-mediated truncation of their translational activity.

**Figure 7.**
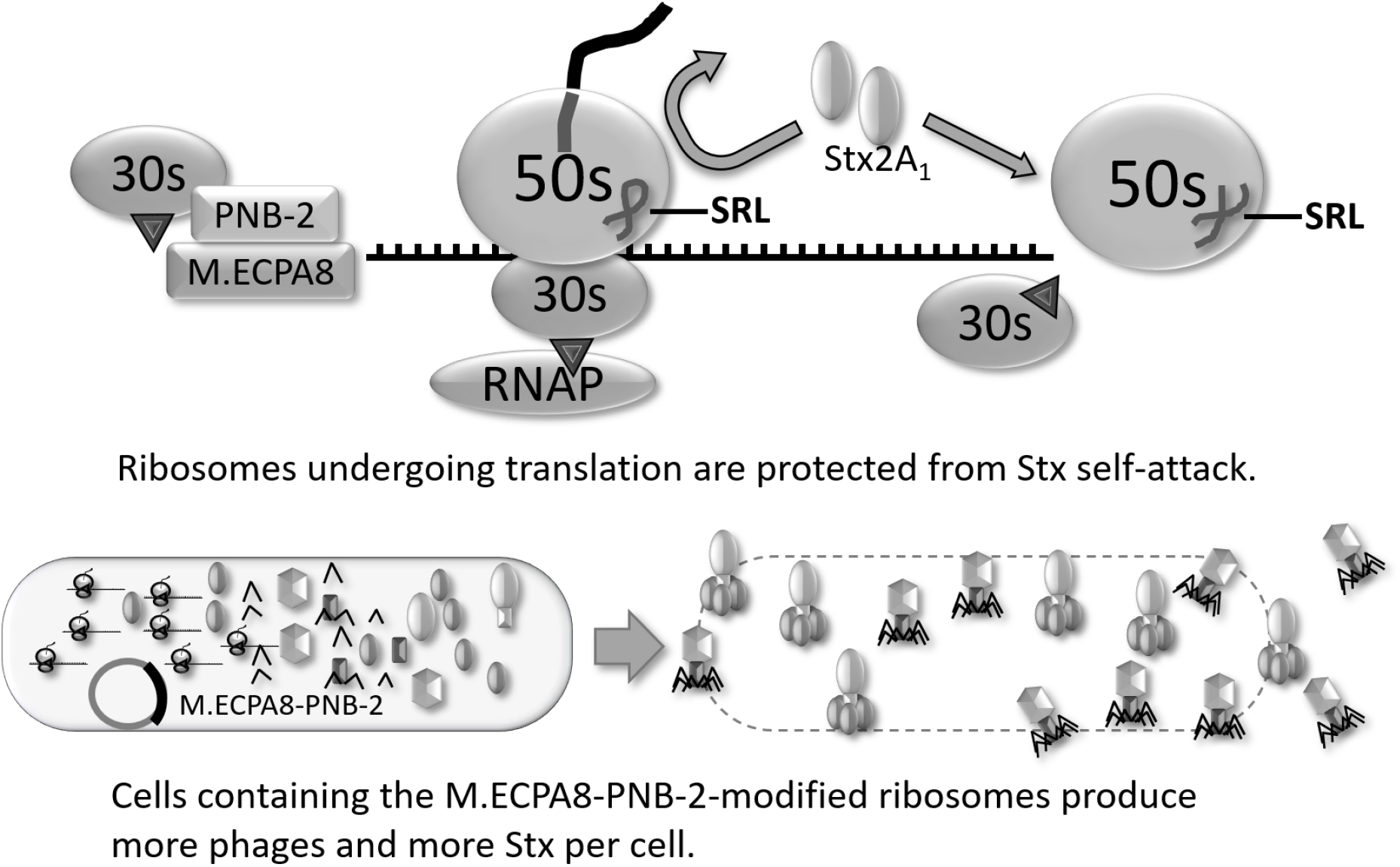
The model of M.ECPA8_3172P-PNB-2 regulates prophage spontaneous induction. The methylation of 16s rRNA performed by M.ECPA8_3172P-PNB-2 inhibit Stx-mediated ribosome inactivation during the translation. Once translation finishes, the ribosome dissociates into 30s and 50s ribosomal subunits. Then the Stx can attack the free 50S subunit prior to its reassembly into a new expressome and trigger cell lysis. The STEC strain with M.ECPA8_3172P-PNB-2-modified ribosomes has higher translation efficiency and produce more phages as well as more Stx per cell, enhancing virulence.

Although our model accounts for the effects of M.ECPA8_3172P-PNB-2 on virulence and phage production, one question remains at issue; how does methylation of the small subunit rRNA prevent Stx-mediated ribosome inactivation? This question is especially puzzling since the 1) Stx binding site is located on the P-stalk region, which is on the large subunit (*13, 56–61*); 2) Stx acts by removing an adenine base from sarcin-ricin loop (SRL) present in 23S rRNA; 3) the putative positions of the bases modified by M.ECPA8_3172P- PNB-2, are on a solvent exposed surface of 30S subunit (Figure 6b & c), that is on the opposite face of the ribosome from the SRL and the P stalk (data not shown). Hence it seems unlikely that methylation directly inhibits Stx binding to the ribosome.

It is possible that the translation enhancement provided by M.ECPA8_3172P-PNB-2 methylation in Stx-encoding cells may be a consequence of increased rates of ribosome maturation. According to this idea, methylation would have the effect of increasing and/or maintaining the number of active ribosomes, “diluting out” the effect of Stx. However, although we cannot rule out an effect of methylation on ribosome maturation, the finding that methylation inhibits Stx-mediated ribosome inactivation *in vitro* (Figure 5) argues against this alternative.

Alternatively, the observation that the putative methylated bases are located on the surface of the ribosome that interacts with the transcribing RNA polymerase (Figure 6d (*49–52*)) suggests that the translation enhancement provided by M.ECPA8_3172P-PNB-2 methylation in Stx-encoding cells is related to transcription-translation coupling. If this inference is correct, we imagine that M.ECPA8_3172P-PNB-2 methylation may enhance transcription-translation coupling. As a consequence, the steady-state occupancy of the translating ribosome with EF-Tu and EF-G would be increased. The SRL is part of the binding site for these factors on the ribosome. Therefore, we suggest that the increased steady-state occupancy of EF-Tu and/or EF-G could block Stx from accessing its substrate in the SRL. That is, methylation may indirectly block Stx-mediated ribosome inactivation by stabilizing the expressome. This suggestion is consistent with the finding that methylation of 16s rRNA cannot completely block Stx-mediated ribosome inactivation in our *in vitro* transcription system. That is, after finishing translation, the ribosome dissociates into its component subunits and we envision that as the Stx concentration increases, it can attack a greater fraction of the 50S subunit prior to its reassembly into a new expressome. This view of M.ECPA8_3172P-PNB-2 mechanism of action indicates that its role is merely to delay, rather than completely prevent, ribosome inactivation. However, our results indicate that this delay plays a critical role, allowing the Stx-encoding phage to accumulate more Stx and phage particles before cell lysis, thereby completing a more productive infectious cycle and in so doing increase its fitness, while also increasing the virulence of its bacterial host. Overall, the findings suggest the Stx-encoding prophage is not merely a toxin gene carrier, instead it actively modulates the production and releasing of the carried toxin to affect the feature of its bacterial host.

## Methods and Materials

### Strains

*Acanthamoeba castellanii* strain (*A*. *castellanii* ATCC 30234) and *E coli.* MG1655 strain were obtained from the American Type Culture Collection (Manassas, VA). PA2 and PA8 phages were obtained by inducing O157:H7::ϕPA2 and O157:H7::ϕPA8 strain with 50 μg/ml ciprofloxacin for 8h. *E*. *coli* O157:H7 strains containing PA2 or PA8 were a generous gift from Dr. Edward G Dudley, Penn State University. Each strain harbors only one Stx-encoding prophage and those prophages encode *stx*2a. These genomes of these strains and the isolated phages have been sequenced (GenBank accession numbers: *Escherichia coli* PA2 NZ_AOEL00000000; *Escherichia coli* PA8 NZ_AOEO00000000 ; ϕPA2 KP682371.1; ϕPA8 KP682374 (*62*).

PA2 and PA8 phages were obtained by inducing *Escherichia coli* PA2 or *Escherichia coli* PA8 strains with 50 μg/ml ciprofloxacin for 8h. Respective lysogens in MG1655 were constructed as described (*63, 64*). Briefly, dilutions of phages isolated from the parental EHEC strains were spotted on a lawn of MG1655 growing in soft agar overlaid on an LB agar plate and the plates incubated overnight at 37 °C to allow the formation of phage plaques. Putative lysogens were picked from the center of these plaques and streaked on an LB agar plate. The successful lysogenization was verified by colony PCR.

### Growth conditions

All the overnight bacterial culture used in experiment were grown from single colonies in Luria-Bertani broth (LB) medium supplemented 10 mM MgSO_4_ at 30 °C. SOC medium (*65*) was used in the recovery of the electro-competent cells. LB medium supplemented with chloramphenicol (Cm) (10 μg/ml or 20 μg/ml) and ampicillin (Amp) (100 μg/ml) was used in the selection of knockout lysogens.

### Plasmids

Plasmids were purified using the Qiagen Plasmid Midi or Max purification kit (Qiagen Inc., Valencia, USA). pTXB1-*sfGFP* plasmid (p-*GFP*) used in CFPS to produce GFP was purchased from Addgene Inc. (MA, USA). The p-*stx_2_* plasmid encoding Stx2 holotoxin was constructed and tested as described (*66*). pKD3 was obtained from Addgene. pKD46*recx* was a gift from Michael Berger, University of Műnster.

### Knock-out phages

Recombination substrates used to replace the *gp5* or *gp6* that encodes PNB-2 or M.ECPA8_3172P-PNB-2 in MG1655::ϕPA8 with a chloramphenicol acetyl transferase gene (*cat*), were generated using high-fidelity Platinum *PFX* DNA polymerase (Invitrogen, Thermo-Fisher Inc., Waltham, MA) using pKD3 as templates with following primers sets: PNB-2F and PNB-2R (TM 62°C) for *gp6* gene, PNB-2F and M.ECPA8_3172PR for both *gp5* and *gp6* genes (Table 1). The knockout prophage was constructed using a lambda red system as described (*67*) with modifications, specifically; pKD46*recx* plasmid was used instead of pKD46 (*68*); 2, electro-transformation was performed using a Bio-Rad Gene pulser with following setting: 2.5kV, 25F, 200 ohm and 5ms; 3, primers Def1F and DetR were used to screen presumptive recombinants (Table 1). The mutated phages were induced with 50 μg/ml ciprofloxacin for 8h and the isolated phage were used to formed lysogens in *E. coli* MG1655 strain as described above.

**Table 1.**
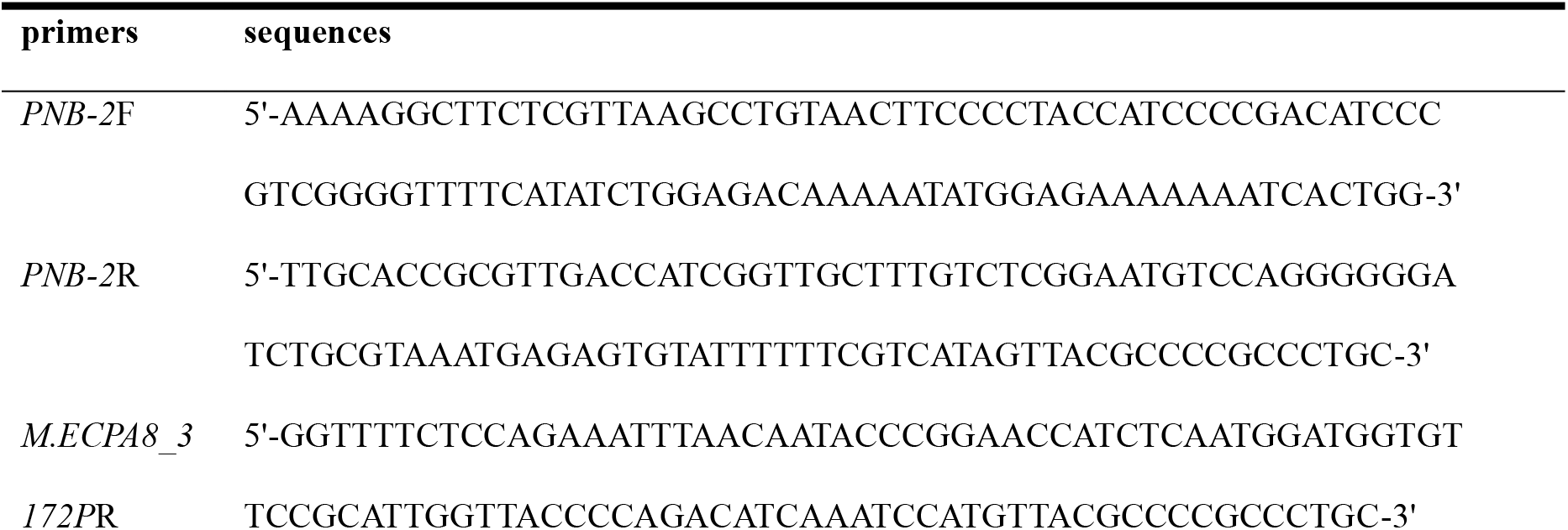
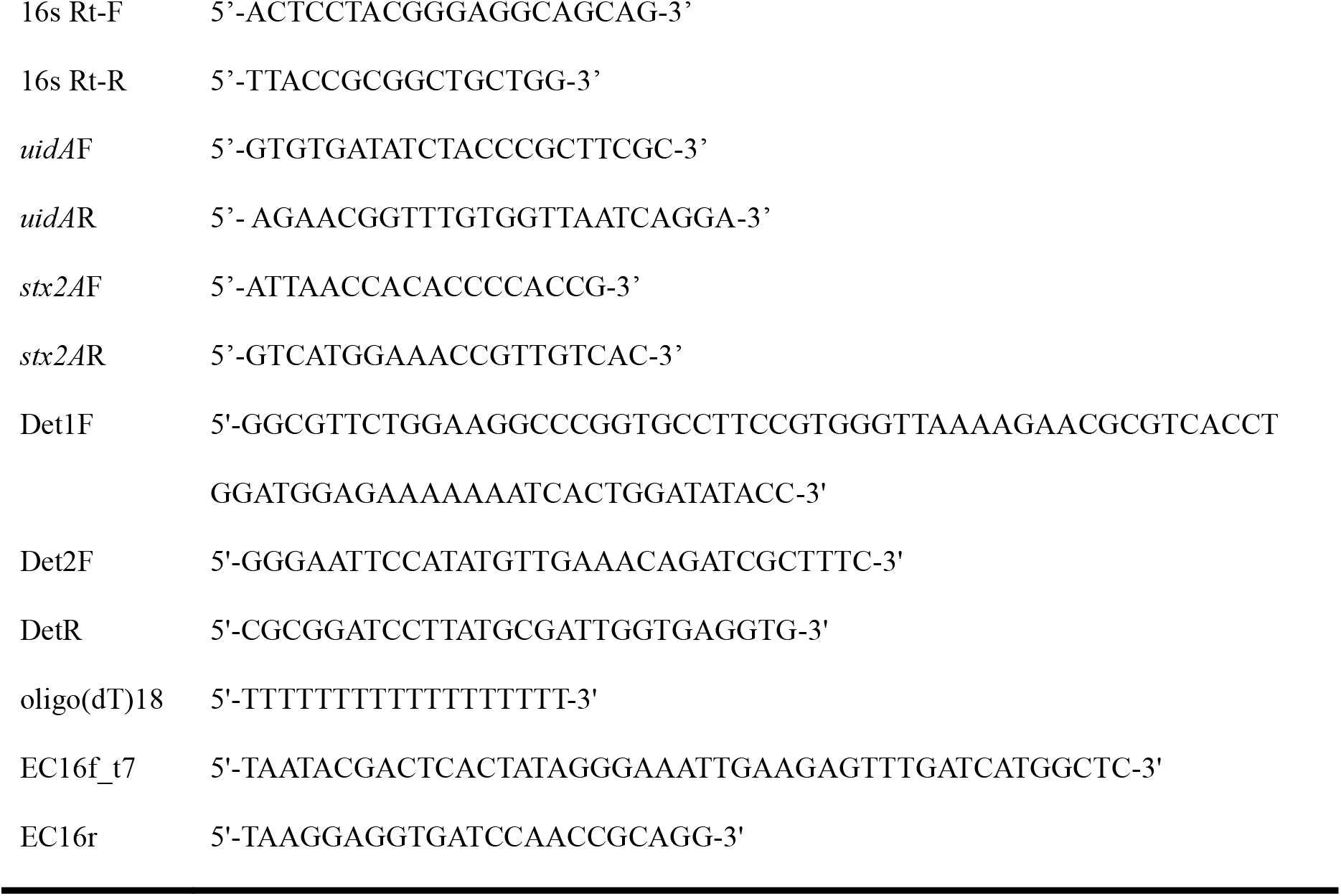
Oligonucleotides used in this study.

### Acanthamoeba Predation assay

The predation assay was performed essentially as described previously (*69*). Briefly, proteose peptone glucose agar plates (1% proteose peptone, 1% glucose) were separately seeded with 100 μl of saturated cultures (10^9^ cells/ml) of different *E. coli* strains and incubated for 1 hour at 37℃. Subsequently, 10^5^ *Acanthamoeba. castellanii* trophozoites in 10 μl were spotted on each plate. Each plate was sealed with parafilm and incubated at 28.5℃ for 5 days. After incubation, the plates were photographed and the size of plaque formed by *A. castellanii* consumption on each plate was calculated using ImageJ (*70*). Each measurement was done in quadruplicate and averaged. The data shown represent the averaged results of ≥ 3 independent quadruplicate measurements.

### Effect of M.ECPA8_3172P-PNB-2 deletion on transcription of downstream genes

Overnight cultures of MG1655::ϕPA8 and MG1655::ϕPA8Δ*M.ECPA8_3172P*Δ*PNB-2* strains were prepared as described above and the cells were collected by centrifugation at 3000 x *g* for 10 min. Total RNA was extracted using hot acid phenol method (*71*) with two modifications: 1) the 65 °C incubation was for 10 min; 2) the extracted RNA was a chloroform extracted before precipitation. Residual DNA was removed by RNase-free RQ1 DNaseI (Promega, Madison, WI) following the manufacturer’s instructions. The primers set Det2F and DetR was used to create the cDNA from the purified total RNA. RT-PCR was performed with RevertAid reverse transcriptase (Thermo-Fisher, Waltham, MA) following company’s procedure and a no reverse transcriptase control was set up to rule out DNA contamination. The cDNA was examined by regular PCR with the same primer set (TM 60°C) and products were displayed on agarose gel.

### Measurement of Spontaneous Induction and Ribosome abundance

Overnight cultures were prepared as described above. The cells were collected by centrifugation at 3000 x *g*, washed three times with equal volume fresh LB with 10mM MgSO4 and resuspended in equal volume of LB with 10mM MgSO4. Control experiments revealed that washing reduced the amount of free phage to undetectable levels. The washed cells were diluted 1:40 into fresh LB with 10mM MgSO_4_ and incubated at 37°C for 2 h. This time point was determined as time 0. Following this, 100 µl of cultures were taken every 30 min as samples.

Total DNA was extracted from each sample by adding an equal volume of Instagene matrix (BioRad, Carlsbad, CA) The isolated DNA was used as template for qPCR to quantify the amount of phage and bacterial DNA in each sample at various time points. The primers to quantify phage DNA were directed against *stx2A*: *stx2A*F *and stx2A*R (Table 1) and *stx2A* gene probe (*Taq*Man probe 5’-CAGTTATTTTGCTGTGGATATACGA-3’ labeled with fluorescent reporter dye HEX (hexacholoro-6-carboxyfluorescein) at the 5’ end and with the Black Hole Quencher 1 (BHQ 1) at the 3′ end Integrated DNA Technologies, U.S.). The primers used to quantify bacterial DNA were directed against the *uidA* gene: *uidA*F and *uidA*R (Table 1) and *uidA* gene probe (*Taq*Man probe 5’- TCGGCATCC GGTCAGTGGCAG T-3’ which was labeled with the fluorescent reporter dye FAM (fluorescein) at the 5′ end and with BHQ 1 at the 3′ end). The qPCR reaction was performed with Taq DNA polymerase (New England Biolabs, Beverly, MA) according to the manufacturer’s instructions. 0.375 µM of each primer plus 0.3 µM probe were used in a 20 μl qPCR reaction. The qPCR was conducted on a Bio-Rad MyIQ2 platform using the following program: 5 min at 95°C, 45 repeats of 10 s at 95°C, 45 s at 60°C.

Standard curves for qPCR were generated using RNAse-treated DNA purified by phenol-chloroform extraction from isolated PA8 phage and MG1655. The MG1655::ϕPA8 lysogen was used as the source of phage DNA which was isolated by a modified PEG precipitation from ciprofloxacin-induced cultures (*72*). qPCR results are was expressed as the ratio of total phage DNA:total bacterial DNA. Since the genomic DNA of each MG1655::ϕPAx lysogen contains only one prophage, ratios >1 denote the amount of phage released per bacteria. Each data point was determined from at least 3 biological replicates and each biological replicate was measured from at least 3 technical replicates.

For measuring amount the amount of 16s rRNA, total RNA was extracted from the samples using Direct-zol RNA miniprep kit (Zymo Research, Orange, CA) with modifications: 1) After adding the Tri-reagent solution (Thermo-Fisher, Waltham, MA), samples were incubated for 15 min at room temperature; 2) Instead of on-column digestion, residual DNA was removed by RNase-free RQ1 DNaseI (Promega, Madison, WI) following the manufacturer’s instructions. Random hexamers (Thermo-Fisher, Waltham, MA) were used to create the cDNA library from total RNA with RevertAid reverse transcriptase (Thermo-Fisher, Waltham, MA) following the manufacturer’s instructions. The qPCR set-up was performed as described (*73*). Primer set 16s Rt-F and 16s Rt-F was used to quantify 16s rRNA; Primer set *uidA*F and *uidA*R was used to quantify *uidA* mRNA. The qPCR conditions, standard curves generation and statistical analysis of the data were performed as described above.

### Cell-free protein synthesis

S12 extracts were prepared essentially as described (*42*). Briefly, the overnight cultures were prepared as described above. 0.5 mL of the cultures were diluted into 50 mL fresh 2X YTPG (*74*) in a 500 mL flask and incubated at 37°C with shaking for 5h. The cells were collected by centrifugation at 3000 x *g* for 20 min and washed once with S12 lysis buffer (10 mM Tris-acetate pH 7.4, 14 mM magnesium acetate, 60 mM potassium acetate, made with DEPC-treated water) before storing in −80°C as pellets. Subsequently, 0.5 g thawed cells were suspended in 1 ml S12 lysate buffer and lysed by sonication (Sonic Dismembrator Model 500 with 2 mm tip, Thermo-Fisher, Waltham, MA, set to 30% amplitude and 12s/60s on/off pulse) for 8 min. The lysate centrifuged at 12,000 x *g* for 2 min, and the supernatant transferred to fresh tube and centrifuged again for 8 min. The protein concentration in the final supernatant was measured by Bio-Rad protein assay (Bio-Rad, Carlsbad CA) following company’s procedure. The extracts were aliquoted and stored at a protein concentration of 0.1 g/mL in −80°C.

*In vitro* translation reactions were carried out in a 1.7 mL microtube placed on a rotator and maintained at 37°C. The 10 μL reaction consists of the following components: 2.5 μL of S12 extraction, 5 μL of 2X protein synthesis buffer (New England Biolabs, Beverly, MA), 0.25 μL of RNaseOUT RNAse inhibitor (Thermo-Fisher, Waltham, MA), 500 ng of purified p-*GFP* plasmid, 0.1 μL of T7 RNA Polymerase (High Concentration, New England Biolabs, Beverly, MA), 20 mM Tris-acetate pH 8.5 and DEPC-treated water. After 5h incubation, the amount of cell-free synthesized GFP protein was estimated by measuring the florescence intensity using Spark multimode microplate reader (Tecan Trading AG, Switzerland).

To measure the relative translation efficiency of S12 extraction made from different lysogen in presence of Stx2, a modified CFPS system was used. In this case, 22.5 nM, 18 nM, 13.5 nM or 9 nM of either p-*stx_2_* or pET17b (control) were first added into the complete CFPS reaction prior. After 30 min incubation at 37°C, 500 ng of p-*GFP* plasmid was added into the reactions and then incubated for another 2 h before measuring the fluorescence intensity. The effect of p-*stx_2_* on translation of GFP was calculated as percent of control.

### The preparation of the 16S rRNA samples for direct sequencing

16S rRNA was extracted from the purified 30S small ribosome subunits isolated from MG1655 and MG1655::ΦPA8 strain. In brief, 600 ml of OD_600_= 0.6 bacterial culture was harvested by centrifugation at 3,000 × *g* for 15 min and was washed with 25 ml S100 buffer (30 mM Tris pH 7.9, 3 mM MgCl_2_, 140 mM KCl, 6 mM BME) with 200 µl protease inhibitor cocktail and then was resuspended in 25ml S100 buffer. All subsequent procedures were performed at 4°C or on ice. The resuspended cells were lysed by passing through a French press twice as describe above. The lysate was clarified by centrifugation at 15,000 x g for 15 min at 4°C twice. The ribosomes were collected by ultra-centrifugation at 32,700 x g for 4h at 4°C. The collected ribosomes were flash frozen and stored at −80 °C. For use, the ribosome pellet suspended in 5 ml 30-50 Buffer A (50 mM Tris pH 7.0, 10.5 mM MgCl_2_, 100 mM NH_4_Cl, 6 mM BME) and subunits separated by velocity gradient sedimentation on 10-30% sucrose (v/v) gradients by centrifugation at 25,000 *x g* for 14 h at 4°C in a Beckman SW28 rotor. Gradient fractions were separated on Gradient Station spv1.5 (BioComp, Canada) while scanning at 254 nm and subunit containing fractions were pooled and stored at −80 °C. Where desired, rRNA was extracted from the gradient fractions contain 30S or 50S ribosome subunits by a modified phenol/chloroform extraction. Briefly, 1 volume phenol pH 8.0 was added to the fractions then vortexed for 1 min prior to 5 min centrifugation at 14000 *x g*. Subsequently, the samples were extracted with 1 volume of acid phenol/chloroform (5:1) pH 4.5, followed by extraction with 1 volume of chloroform. The extracted rRNA was precipitated by ethanol precipitation (adding 3 volumes of ethanol and 0.1 volume of 3 M sodium acetate and then incubating on dry ice for 1h), centrifuged for 10 minutes at 10,000 × *g*. The pellet was washed with 1 volume of 70% ethanol. The RNA pellet was resuspended in DEPC-treated H_2_O and used directly in the polyA tailing reaction as described below.

An unmethylated 16S rRNA control was made by *in vitro* transcription, Briefly, template DNA used to transcribe 16S rRNA was PCR amplified from genomic DNA of MG1655 strain purified by regular phenol-chloroform extraction and ethanol precipitation using the Taq KeenGreen 2x Master Mix (IBI Scientific, U.S.) and the primer set EC16f_t7 and EC16r (*75*). Subsequently. 30 μL of the PCR product was added directly to a 50 μL *in vitro* transcription reaction with T7 RNA Polymerase (New England Biolabs, Beverly, MA) according to the manufacturer’s instructions and incubated for 3h at 37 ℃. The *in vitro* transcription product was used directly in the polyA tailing reaction as described below.

All the 16S rRNA used for nanopore sequencing were treated with polyA polymerase (New England Biolabs, Beverly, MA) according to the manufacturer’s instructions. In brief, 5 μg of the purified 16S rRNA or 15 μL of the *in vitro* transcription product was used in a 20 μL system and incubated for 45 min at 37 ℃. The products were subsequently treated with RQ1 RNase-free DNase as described by the manufacturer for 60 min to remove template DNA. After adding 3 volumes of Tri reagent and 4 volume ethanol, the RNA samples were purified with Direct-zol RNA Midiprep Kits (Zymo Research, Orange, CA) without the in-column DNA digestion step. The addition of the polyA tail was confirmed by analyzing a portion of the reaction on a MOPS/formaldehyde denaturing agarose gels (*76*) and by a reverse-transcription PCR with oligo(dT)18 primer using SuperScript™ III Reverse Transcriptase (Thermo-Fisher, Waltham, MA) following company’s procedure.

### The sequencing of 16S rRNA

Direct rRNA sequencing was performed using MinION chemistry flowcells according to the instructions provided by Oxford Nanopore Technologies (Oxford, UK). Three different polyA+ rRNA were sequenced using the provided polyT adapter. The methylation positions on the rRNA purified from MG1655 and MG1655::ΦPA8 were determined by pairwise comparison of signal level ionic flow features using the xpore (*48*) procedures against the unmethylated 16S rRNA control synthesized by in vitro transcription, as mentioned above. The dataset used for comparison was cleaned by PHRED to remove reads with a quality score lower than 10 or a basecall accuracy lower than 90%. The cleaned dataset was processed using samtools with the parameter ‘-q 60’ to identify unique reads. These reads were then mapped to the 7 variants of the 16S ribosomal gene using minimap2 software. Results with a read coverage outside the 15-100,000 range were excluded. Gaussian distributions were estimated using Variational Bayesian inference, and modification rates were calculated for each potential modification site accordingly. Next, we performed a z-test between the rRNA purified from M.ECPA8_3172P-PNB-2-replete strain (MG1655::ΦPA8) and the rRNA purified from the MG1655 strain to obtain the Benjamini-Hochberg adjusted p-values for significance. Meanwhile, Wilcoxon rank-sum test was implemented to remove background noise. Potential methylation sites of M.ECPA8_3172PPNB-2 were identified based on a corresponding log_10_(p-value) smaller than −100 and a change in RNA modification rate greater than 0.10.

### Statistical Analysis

Error bars in the figures represent standard deviations of multiple replicates experiments. All the comparison between two groups were analyzed by unpaired T-test using GraphPad PRISM (version 4.0). For more than two groups, nonpaired one-way analysis of variance (One-Way ANOVA) using GraphPad PRISM (version 4.0) followed by Tukey’s multiple comparisons test to determine the statistical significance between different samples was used. All the Measurements were made from at least 3 biological replicates and each biological replicate was measured from at least 3 technical replicates.

