## Supplemental Figure 1 for "The Shiga toxin (Stx)-Phage Encoded Ribosomal RNA Methyltransferase Regulates Stx-producing *Escherichia coli* (STEC) Virulence by Blocking Stx-Mediated Inactivation of Bacterial Ribosomes"

### Supplementary Information

#### Supplementary Figures

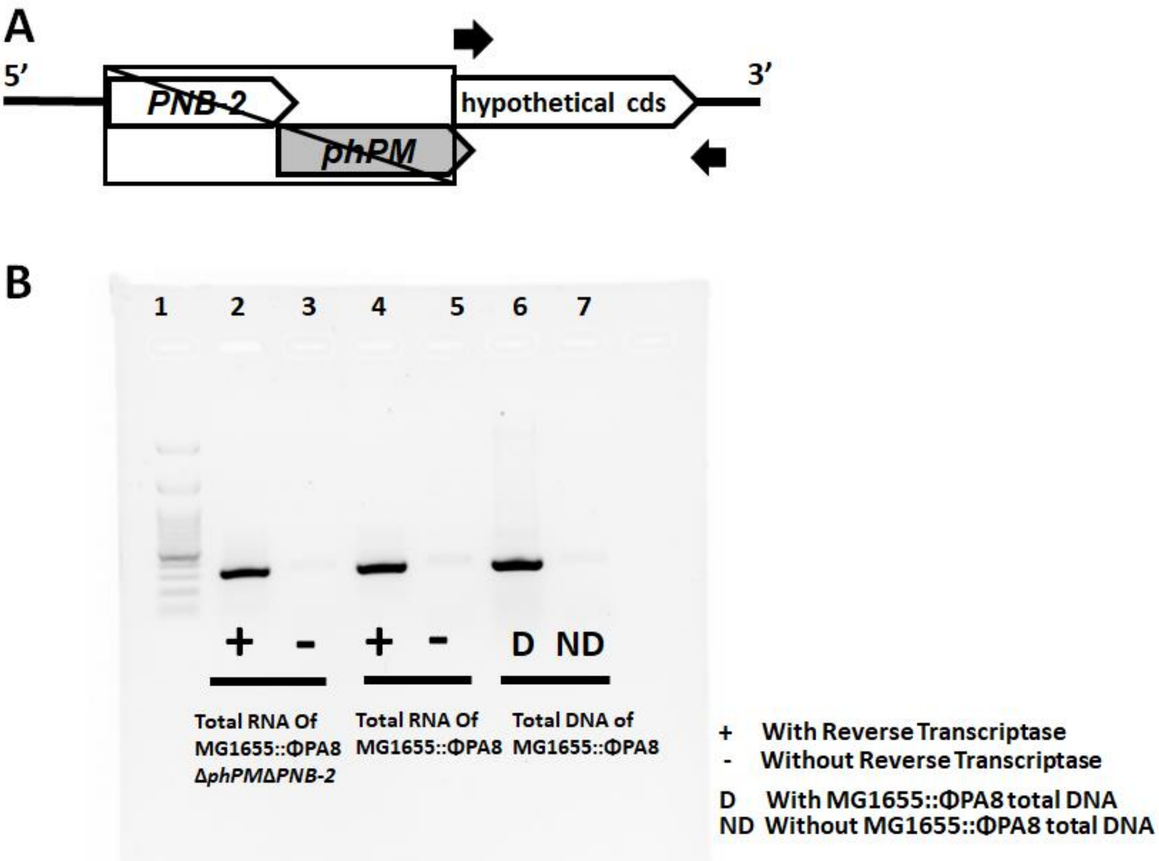


**Figure S1. The transcription of *M.ECPA8_3172P* downstream gene in double-knockout strain is unaffected.** The total RNA that extracted from the MG1655::ϕPA8 and MG1655::ϕPA8Δ*M.ECPA8_3172P*Δ*PNB-2* strain was reverse-transcribed into cDNA and checked as described in Methods and Materials. **A.** The cDNA was checked by regular PCR with two primers that amplify the downstream hypothetical coding region of *M.ECPA8_3172P* (primers were indicated by the arrows). **B.** The PCR product was displayed by ethidium bromide staining on an agarose gel. The bands in lane 2&4 represent the cDNA that synthesized from the mRNA molecule purified from double-knockout strain and wild-type strain. The lane 3&5 are PCR reactions using total RNA that has not been subjected to reverse transcription. Lane 6&7 displays the PCR product obtained using total DNA from MG1655::ϕPA8 or $ddH_2O$ as template.
